# Improvement of muscle strength in a mouse model for congenital myopathy treated with HDAC and DNA methyltransferase inhibitors

**DOI:** 10.1101/2021.11.09.467894

**Authors:** Alexis Ruiz, Sofia Benucci, Urs Duthaler, Christoph Bachmann, Martina Franchini, Faiza Noreen, Laura Pietrangelo, Feliciano Protasi, Susan Treves, Francesco Zorzato

**Affiliations:** Neuromuscular Research Group, Department of Biomedicine, Basel University Hospital, Hebelstrasse 20, 4031 Basel, Switzerland; Division of Clinical Pharmacology & Toxicology, Department of Biomedicine, University and University Hospital Basel, Switzerland; Genome plasticity group, Department of Biomedicine, University of Basel, Mattenstrasse 28, Basel, Switzerland; CAST, Center for Advanced Studies and Technology & DMSI, Dept. of Neuroscience, Imaging and Clinical Sciences, Univ. G. d’Annunzio, 66100 Chieti, Italy; Department of Life Science and Biotechnology, University of Ferrara, Via Borsari 46, 44100, Ferrara, Italy

## Abstract

To date there are no therapies for patients with congenital myopathies, muscle disorders causing poor quality of life of affected individuals. In approximately 30% of the cases, patients with congenital myopathies carry either dominant or recessive mutations in the *RYR1* gene; recessive *RYR1* mutations are accompanied by reduction of RyR1 expression and content in skeletal muscles and are associated with fiber hypotrophy and muscle weakness. Importantly, muscles of patients with recessive *RYR1* mutations exhibit increased content of class II histone de-acetylases and of DNA genomic methylation. We recently created a mouse model knocked-in for the p.Q1970fsX16+p.A4329D RyR1 mutations, which are isogenic to those carried by a severely affected child suffering from a recessive form of RyR1-related multi-mini core disease. The phenotype of the RyR1 mutant mice recapitulates many aspects of the clinical picture of patients carrying recessive *RYR1* mutations. We treated the compound heterozygous mice with a combination of two drugs targeting DNA methylases and class II histone de-acetylases. Here we show that treatment of the mutant mice with drugs targeting epigenetic enzymes improves muscle strength, RyR1 protein content and muscle ultrastructure. This study provides proof of concept for the pharmacological treatment of patients with congenital myopathies linked to recessive *RYR1* mutations.

## Introduction

Skeletal muscle contraction is initiated by a massive release of Ca^2+^ from the sarcoplasmic reticulum (SR) via the opening of the ryanodine receptor 1 (RyR1), a calcium release channel, which is localized in the SR terminal cisternae (1–3). The signal causing the opening of the RyR1 is the depolarization of the sarcolemmal membrane, which is sensed by voltage-dependent L-type Ca^2+^ channels (dihydropyridine receptor, DHPR) located in invaginations of the sarcolemma referred to as transverse tubules (TTs)(1, 4). Communication between DHPR and RyR1 occurs in specialized in tricellular junction between TT and SR called Ca^2+^ release units (CRUs). Skeletal muscle relaxation is brought about by SR Ca^2+^ uptake via the activity of the sarco(endo)plasmic reticulum CaATPAses (SERCA)(5). Dis-regulation of Ca^2+^ signals due to defects in key proteins (RyR1 and DHPR) involved in excitation-contraction (EC) coupling is the underlying feature of several neuromuscular disorders (6–8). In particular, mutations in *RYR1*, the gene encoding RyR1, are causative of malignant hyperthermia (MH; MIM #145600), central core disease (CCD; MIM #11700), specific forms of multi-minicore disease (MmD; MIM # 255320) and centronuclear myopathy (CNM)(6–9). A great deal of data has shown that *RYR1* mutations result mainly in four types of channel defects (7). One class of mutations (dominant, MH-associated) causes the channels to become hypersensitive to activation by electrical and pharmacological stimuli (9). The second class of *RYR1* mutations (dominant, CCD-associated) results in leaky channels leading to depletion of Ca^2+^ from SR stores (7, 8). A third class of *RYR1* mutations also linked to CCD causes EC uncoupling, whereby activation of the voltage sensor Cav1.1 is unable to cause release of Ca^2+^ from the SR (10). The fourth class comprises recessive mutations, which are accompanied by a decreased content of mutant RyR1 channels on SR membranes (11–14).

Patients with Congenital Myopathies such as MmD carrying recessive *RYR1* mutations belonging to class 4 channel defects, typically exhibit non-progressive proximal muscle weakness (15, 16). This reduced muscle strength is consistent with the lower RyR1 content observed in adult muscle fibers that should result in a decrease of Ca^2+^ release from the SR (11–15). The decrease of RyR1 expression is also associated with moderate fiber atrophy, which may additionally contribute to the decrease of muscle strength. In addition to the depletion of RyR1 protein, muscles of patients with recessive *RYR1* mutations exhibit striking epigenetic changes, including altered expression of microRNAs, an increased content of HDAC-4 and HDAC-5 and hypermethylation of more than 3600 CpG genomic sites (17–19). Importantly, in muscle biopsies from 4 patients hypermethylation of one of the internal *RYR1* CpG islands correlated with the increased levels of HDAC-4 and HDAC-5 (18).

In order to study in more detail the mechanism of disease of recessive *RYR1* mutations, we developed a mouse model knocked in for two mutations, isogenic to those identified in a severely affected child with recessively inherited MmD (16). Such a mouse model carries the p.Q1970fsX16+p.A4329D RyR1 mutations (20) and will henceforth be referred to as double heterozygous (dHT). Characterization of the muscle phenotype of dHT mice demonstrated that it faithfully recapitulates not only the physiological and biochemical changes, but also the major muscle epigenetic signatures observed in muscle biopsies from MmD patients. In the present study we treated dHT mice with drugs targeting epigenetic enzymes and evaluated the physiological effects of the treatment on muscle function as well as on muscle structure. Our results show that treatment of dHT mice with drugs targeting epigenetic enzymes rescues muscle strength, increases RyR1 protein content and improves muscle morphology, i.e. the treatment partially rescues Ca^2+^ release units (CRUs, the intracellular sites containing RyR1), and mitochondria. This study provides proof of concept for the treatment of patients with congenital myopathies linked to recessive *RYR1* mutations, with small molecules inhibiting DNA methyltransferases and histone deacetylases.

## Results

### Effect of TMP269 and 5-Aza-2-deoxycytidine on the *in vivo* muscle phenotype of dHT mice

We first examined the pharmacokinetics and bio-distribution of TMP269, the class IIa-HDAC inhibitor we selected for this study. After intraperitoneal injection of 25 mg/kg body weight of TMP269 dissolved in Polyethylenglycol 300 (PEG300) (500 μl/Kg) and N-Methyl-2-pyrrolidone (NMP) (250 μL/Kg), blood and/or skeletal muscles were collected at different time points and the content of TMP269 was quantified by liquid chromatography tandem mass spectrometry (21). The peak blood concentration of TMP269 was achieved approximately 1 hour after injection. The circulating levels of TMP269 decay within 12 hours (Supplementary Fig. S1A). Importantly, the class II-HDAC inhibitor diffuses into skeletal muscle (Supplementary Fig. S1B), and as expected, its concentration profile in skeletal muscle follows that observed in blood. Although the level of TMP269 accumulating in skeletal muscle is lower compared to that present in blood, its concentration in muscle is adequate to induce an inhibitory effect on class IIa HDACs activity (22). An identical protocol was used to monitor the optimal dose of 5-Aza-2-deoxycytidine (5-Aza), an FDA approved DNA methyltransferase (DNMT) inhibitor (23). The administration of TMP269+5-Aza for 15 weeks resulted in hypomethylation of 165 protein-coding genes (Supplementary Table S1) in soleus muscles. Additionally, administration of TMP269+5-Aza for 15 weeks increases the acetylation of Lys residues (Supplementary Fig. S2A) and of H3K9 (Supplementary Fig. S2B and S2D) in total homogenates from FDB fibers isolated from WT and dHT mice, compared to that observed in fibers from vehicle treated WT and dHT mice.

Having ascertained that the drugs reached and exerted their activity on skeletal muscles we injected 6 weeks old WT and dHT mice with vehicle alone (PEG300+ NMP), with TMP269 alone (25 mg/kg), 5-Aza alone (0.05 mg/kg) or with the two drugs combined on a daily basis and investigated their effects on the *in vivo* skeletal muscle phenotype by analyzing forelimb grip force using a grip strength meter. Administration of each drug singly, namely TMP269 or 5-Aza alone, does not induce any change in the grip strength of WT (Fig. 1A top left and middle panels). The combined drug treatment did not affect the grip strength of WT mice (Fig. 1A, top right panel). In dHT mice, TMP269 alone causes a small but significant increase of grip strength after 10 weeks of treatment (Fig1A low left panel). However, the increased grip strength in dHT mice (Fig. 1A lower right panel) was more evident by the combined drug treatment. In particular, this effect became apparent 4/5 weeks after starting the drug treatment and peaked at 10 weeks. The combined drug treatment rescues approximately 20% of muscle grip strength in dHT mice. Based on these results, we performed all subsequent experiments only with the combined drug treatment (i.e. TMP269+5-Aza).

**Figure 1:**
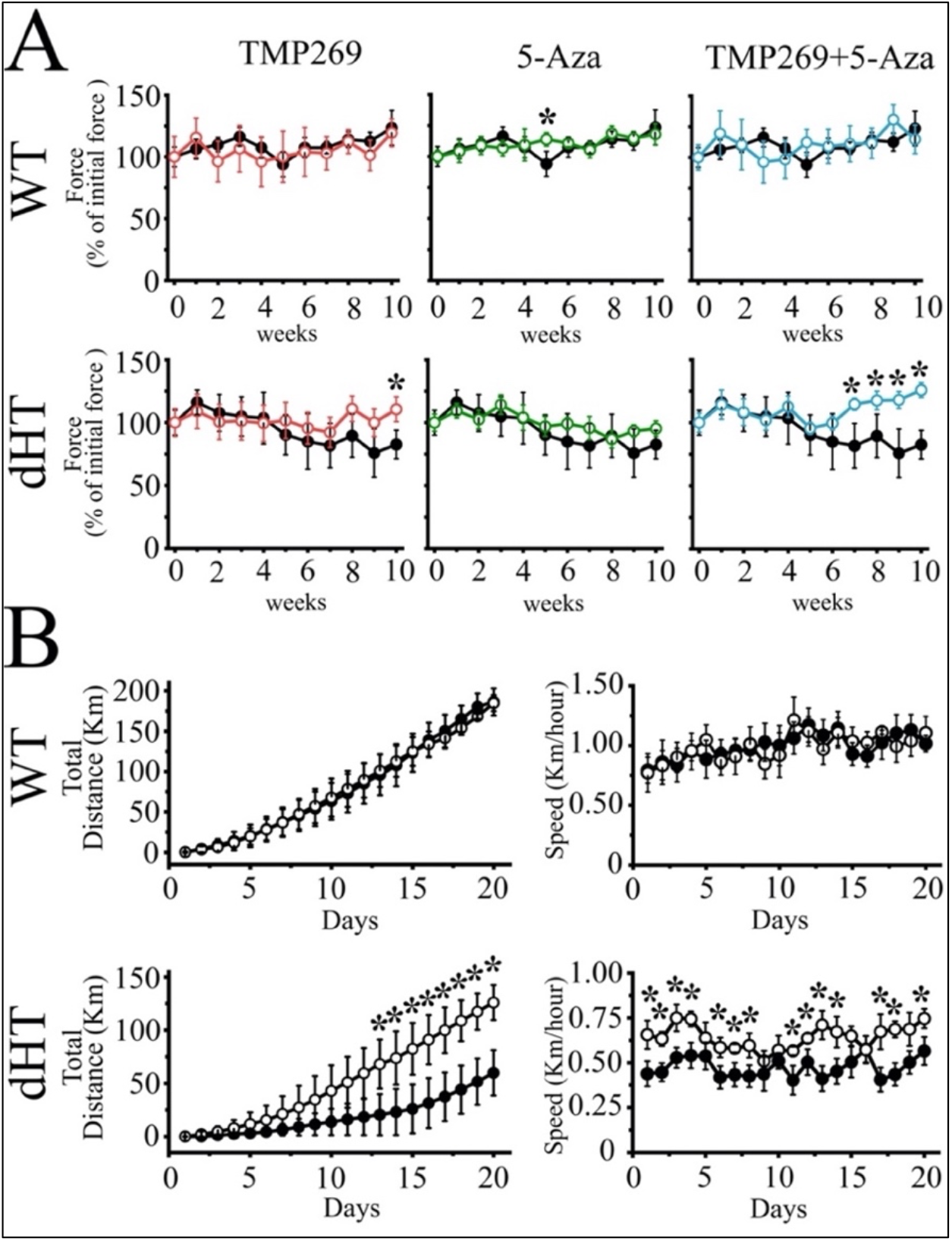
Treatment of dHT mice with TMP269+5Aza improves *in vivo* muscle function as assessed using the grip strength test and voluntary running wheel. **A.** Forelimb (2 paws) grip force measurement of WT (upper panels) and dHT (lower panels) mice treated with vehicle (WT, n= 9; dHT, n=10), TMP269 (WT, n= 11; dHT, n=6), 5-Aza (WT, n= 5, dHT, n=10) and TMP269+5-Aza (WT, n= 10, dHT, n=13). Grip strength was performed once per week during a period of 10 weeks. Each symbol represents the average (±S.D.) grip force obtained in the indicated number (n) of mice. Grip force (Force) values obtained on the first week were considered 100%. Black symbols, vehicle treated-mice, colored symbols, drug-treated mice. Statistical analysis was conducted using the Mann–Whitney test. *p<0.05. **B.** Spontaneous locomotor (dark phase) activity (left panel) and total running speed (right panel) measured over 20 days in 21-week-old dHT and WT littermates mice treated with vehicle or TMP269+5Aza. Data points are expressed as mean (±S.D; n=4-5 individual mice). *p < 0.05 (Mann– Whitney test). The exact P value for day 20 is given in the text.

We next assessed *in vivo* muscle function of WT and dHT vehicle treated or drug treated mice using the voluntary running wheel. We calculated the total running distance of WT (Fig.1B, top panels) and dHT (Fig.1B, bottom panels) mice injected with vehicle and compared it to that of mice treated for 15-18 weeks with TMP269+5-Aza. Three weeks of training improved running performance in both mouse groups. Nevertheless, on day 20 the total running distance of WT mice injected with vehicle alone was approx. 70 % greater compared to that of vehicle treated dHT mice: the total running distance of vehicle treated WT and dHT mice was 186.27±16.70 km n=4 vs 59.93±21.38 km n=5, respectively (mean±S.D., Mann-Whitney two-tailed test, calculated over the 20 days of running, *p<0.05). On the other hand, treatment of dHT mice with TMP269+5-Aza has a remarkable effect on the total running distance (Fig. 1B, lower panel). The beneficial effect begins one week after treatment commencement, and on day 20 the total running distance achieved by dHT mice injected with TMP269+5-Aza was two times higher compared to that covered by dHT mice injected with vehicle alone (Fig. 1B lower panel; Mann-Whitney two-tailed test; on day 20, P=0.041): the total running distance for vehicle treated and drug treated dHT mice was 59.93±21.38 km n=5 and 125.90±16.39 km n=5, respectively (mean±S.D., Mann-Whitney two-tailed test, calculated over the 20 days of running *p<0.05). The longer running distance was also associated with an increase median cruise speed of the drug treated dHT mice compared to vehicle-treated dHT mice (Fig. 1B, right panels) (Mann-Whitney two-tailed test, calculated over the 20 days running period, *p<0.05). The epigenetic modifying drugs most likely affect a number of genes, which in turn leads to an improvement of the *in vivo* muscle performance of the dHT mice; the latter effect may result from an improvement of the mechanical properties of skeletal muscles and/or by an influence of the drugs on the metabolic pathways of muscles. In the next set of experiments, we investigated the mechanical properties of intact EDL and soleus muscles from WT and dHT mice after injection of vehicle and of TMP269+5-Aza.

### Effect of TMP269+5-Aza treatment on isometric force development in muscles from WT and dHT heterozygous mice

EDL and soleus muscles isolated from mice treated for 15 weeks with vehicle or TMP269+5-Aza, were stimulated with a single 15 V pulse of 1.0 msec duration a (Fig. 2A, C, E and G) or by a train of pulses delivered at 150 Hz for 400 msec (EDL, Fig. 2B and D) or 120 Hz for 1100 msec (soleus, Fig. 2F and H) to obtain maximal tetanic contracture. The averaged specific twitch peak force induced by a single action potential in EDL from dHT mice injected with vehicle alone was approximately 37% of that obtained from EDLs from WT mice (64.92±13.93 mN/mm^2^, n=10 vs 171.24±29.32** mN/mm^2^, n=8, respectively; mean± S.D.; ANOVA followed by the Bonferroni post hoc test **p<0.01. Supplementary Table S2). The peak force developed after twitch stimulation of soleus muscles from dHT mice injected with vehicle alone was approx. 67% of that of obtained in soleus muscles from WT littermates injected with vehicle (67.55±11.26 mN/mm^2^, n=10 vs 96.58±25.78* mN/mm^2^, n=8, respectively; mean± S.D. ANOVA followed by the Bonferroni post hoc test *p<0.05. Supplementary Table S2). While the combined drug treatment does not affect the force developed by EDL muscles, we found that it induces a 25% increase of the twitch force in soleus muscles from dHT mice compared to that obtained from vehicle treated dHT mice (84.61±14.06 mN/mm^2^, n=13 vs 67.55±11.26* mN/mm^2^, n=10, respectively; mean± S.D. ANOVA followed by the Bonferroni post hoc test *p<0.05. Fig. 2G and Supplementary Table S2). We next examined the force developed during tetanic contractures of EDL and soleus muscles stimulated by a train of pulses delivered at 150 Hz and 120 Hz, respectively. The maximal specific tetanic force developed in EDL muscles from dHT mice injected with vehicle was approximately 20% lower compared to that of EDL muscles from WT mice (373.76±73.16* mN/mm^2^, n=10 vs 452.97±89.59 mN/mm^2^, n=8, respectively; mean±S.D. ANOVA followed by the Bonferroni post hoc test *p <0.05. Fig.2B and D and Supplementary Table S2). We found no effect of the combined drug treatment on the maximal specific force developed by EDL muscles isolated from dHT mice (Supplementary Table S2). The maximal specific tetanic force generation observed in soleus muscles from dHT mice injected with vehicle was 13% lower compared to WT (276.29±40.04* mN/mm^2^, n=10 vs 315.86±56.96 mN/mm^2^, n=8, mean±S.D. ANOVA followed by the Bonferroni post hoc test *p<0.05) (Fig. 2F and H). Contrary to what we observed in EDL muscles, the combined drug treatment fully rescues the maximal tetanic force of slow twitch muscles. Indeed, soleus from dHT mice treated with TMP269+5-Aza for 15 weeks displayed a maximal tetanic force which was 18% higher compared to that of soleus from dHT mice injected with vehicle (334.78±65.74* mN/mm^2^, n=13 vs 276.29±40.04 mN/mm^2^, n=10, mean±S.D. ANOVA followed by the Bonferroni post hoc test * p<0.05) (Supplementary Table S2).

**Figure 2.**
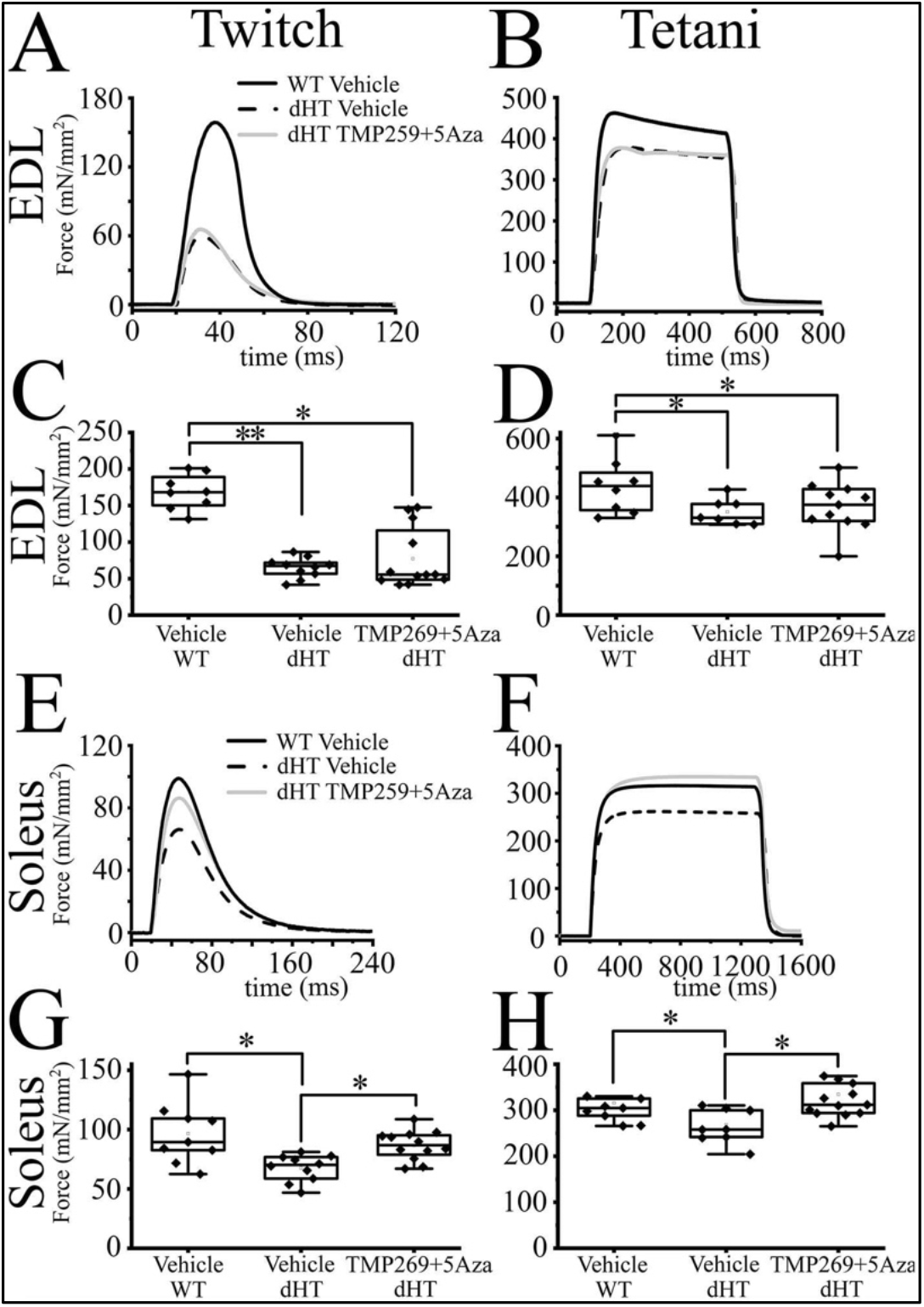
The mechanical properties of dHT treated with TMP269+5-aza improve after 15 weeks of treatment. Mechanical properties of EDL and soleus muscle from WT and dHT mice treated with vehicle (WT, n= 8; dHT, n=10; dHT TMP269+5-Aza, n=13). **A.** Representative traces of twitch and **B.** maximal tetanic force in EDL (150 Hz) muscle from WT and dHT. Force is expressed as Specific Force, mN/mm^2^. **C.** Statistical analysis of force generated after twitch and **D.** tetanic stimulation of isolated EDL muscle. Data points are expressed as Whisker plots (n = 8-13 mice). Each symbol represents the value from a muscle from a single mouse. **E.** Representative traces of twitch and **F.** maximal tetanic force (120 Hz) of soleus muscle from WT and dHT mice. **G.** Whisker plots of force generated after twitch and **H.** tetanic stimulation of isolated soleus muscles. Each symbol represents the value from a muscle from a single mouse (n = 8-13 mice). *p<0.05 **p<0.01 (ANOVA followed by the Bonferroni post hoc test). The exact P values are given in Table S2.

### Fiber type composition and minimal Feret’s diameter of soleus muscles from dHT mice treated with TMP269+5-Aza

In this set of experiments, we examined whether the ergogenic effect associated with the inhibition of epigenetic modifying enzymes is linked to fast-to-slow fiber transition and/or to changes of minimal Feret’s diameter. Treatment with TMP269+5-Aza causes no changes in the content of the fiber type composition of soleus muscles (Fig. 3A and B, and Supplementary Table S3). Similarly, the improved specific force cannot be attributed to a major shift of minimal Feret’s fiber diameter distribution (Fig. 3 C).

**Figure 3:**
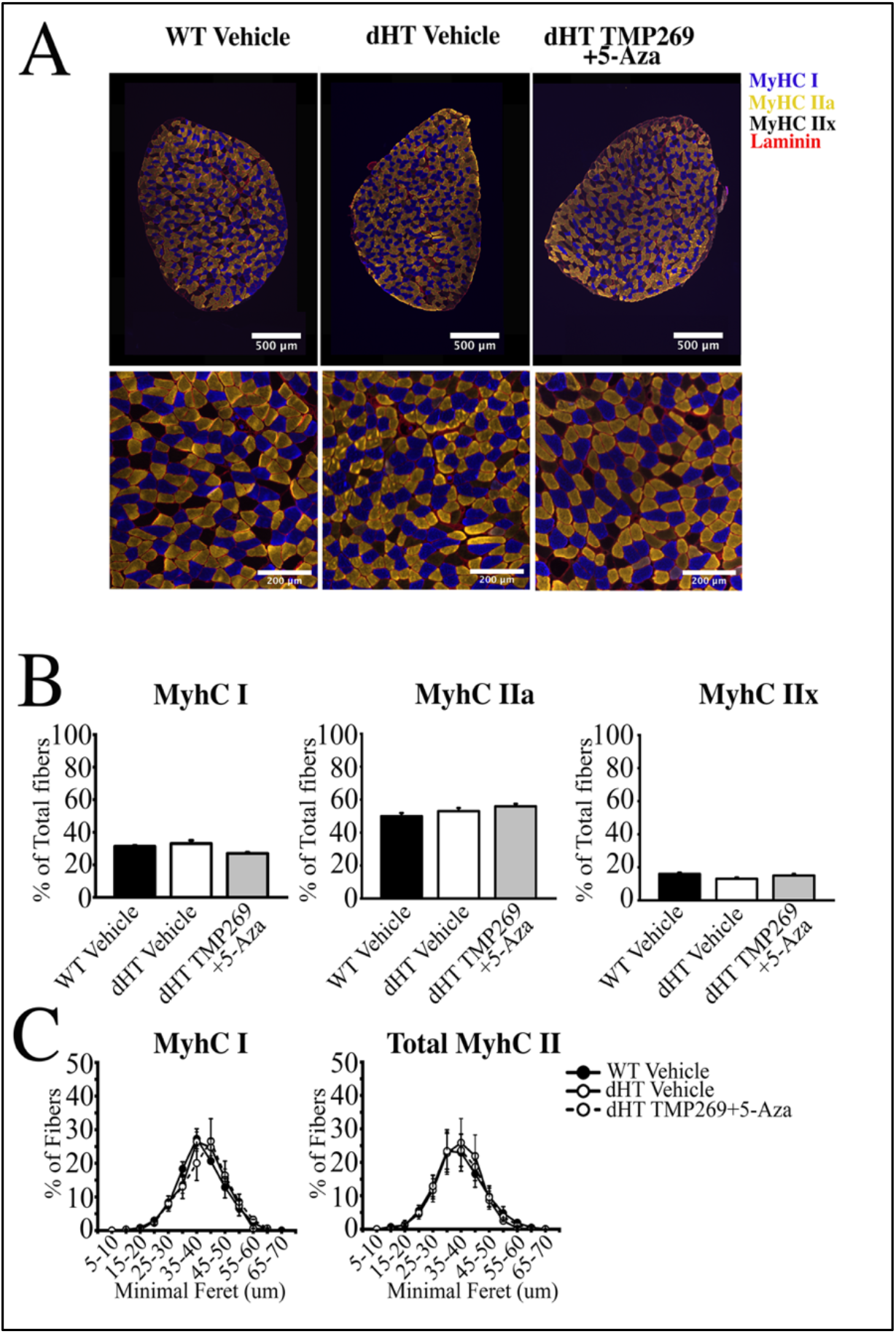
Histology of soleus muscles from TMP269+5-Aza and vehicle treated dHT mice. **A.** Analysis of soleus muscles from WT (vehicle treated) and dHT (vehicle treated and TMP269+5-Aza) mice using monoclonal antibodies specific for myosin heavy chain (MyHC) isoforms. Frozen muscle sections were stained with anti-MyHC I antibodies (slow fibers, blue), anti-MyHC IIa antibodies (fast fibers, yellow) and counterstained with anti-laminin antibodies (red). MyHCIIx (fast fibers) are unstained. **B.** Bar plots of fiber type composition of soleus muscles. Left, mean (%, ± S.E.M.) MyHC I fibers, middle, mean (%, ± S.E.M.) MyHC IIa fibers, right mean (%, ± S.E.M.) MyHC IIx fibers. **C.** Minimal Feret’s distribution of type I and type II fibers. Data points are expressed as mean (± S.E.M.). For WT vehicle treated a total of 2881 fibers from 3 mice were counted, for dHT vehicle treated a total of 2642 fibers from 3 mice were counted, for dHT TMP269+5-Aza a total of 2983 fibers from 3 mice were counted.

### Effect of TMP269+5-Aza treatment on calcium transients in single FDB fibers from WT and dHT mice

We investigated resting [Ca^2+^] and calcium transients evoked either by a single pulse (Fig. 4A and B) or by a train of action potentials (Fig. 4C and D) in *flexor digitorum brevis* (FDB) fibers from WT and dHT mice treated for 15 weeks with vehicle or TMP269+5-Aza (4-6 mice per group). We used single FDB fibers (i) because they are a mixture of fast and slow twitch muscles and (ii) since intact single fibers from EDL and soleus muscles from 22 weeks old mice are nearly impossible to obtain. We found that the resting [Ca^2+^] was similar in FDB fibers from WT and dHT vehicle or drug treated mice. The Fura-2 fluorescence values (F340/F380, mean ±S.D.) were 0.81 ± 0.09 (n= 75 fibers isolated from 4 mice), 0.77 ±0.07 (n= 40 fibers isolated from 5 mice) and 0.81±0.08 (n=33 fibers isolated from 5 mice) in WT mice injected with vehicle, dHT mice injected with vehicle, and dHT mice treated with TMP269+5-Aza, respectively. In the presence of 1.8 mM Ca^2+^ in the extracellular solution, the average peak intracellular Ca^2+^ transient induced by a single action potential in FDB fibers from dHT mice injected with vehicle is approx. 30% lower than that observed in fibers from WT mice (ΔF/Fo values were 0.98± 0.22, n=110 fibers isolated from 6 mice vs 1.38± 0.30, n=91 fibers isolated from 4 mice, respectively; mean± S.D. Fig. 4A and B, Supplementary Table S4). Interestingly, treatment of dHT mice with TMP269+5-Aza causes a 23% increase of the peak Ca^2+^ transient compared to that observed in dHT mice injected with vehicle (*1.21±0.28, n=155 fibers isolated from 5 mice vs 0.98±0.22, n=110 fibers isolated from 6 mice, respectively; ΔF/Fo values are expressed as mean± S.D. ANOVA followed by the Bonferroni post hoc test *p<0.05. Fig. 4A and B and Supplementary Table S4). In the presence of 1.8 mM Ca^2+^ in the extracellular solution, the peak Ca^2+^ transient evoked by a train of pulses delivered at 100Hz in FDB fibers from dHT mice injected with vehicle, is approx. 25% lower compared to that of FDB fibers from WT mice (*1.22±0.22, n=92 fibers isolated from 6 mice vs 1.62±0.21, n=63 fibers isolated from 4 mice, respectively; ΔF/Fo values are expressed as mean± S.D. Fig. 4D and Supplementary Table S4). When dHT mice are treated with TMP269+5-Aza for 15 weeks, the summation of calcium transients induced by a train of supramaximal pulses, is 16% higher compared to that of dHT mice injected with vehicle alone (*1.42±0.23, n=78 fibers isolated from 5 mice vs 1.22±0.22, n=92 fibers isolated from 6 mice, respectively; ΔF/Fo values are expressed as mean± S.D.; ANOVA followed by the Bonferroni post hoc test *p< 0.05. Fig. 4C and D and Supplementary Table S4). Altogether, the increase of the peak calcium transient after either a single action potential or a train of pulses is consistent, at least in part, with the ergogenic effects caused by the combined treatment with TMP269+5-Aza on dHT mice.

**Figure 4.**
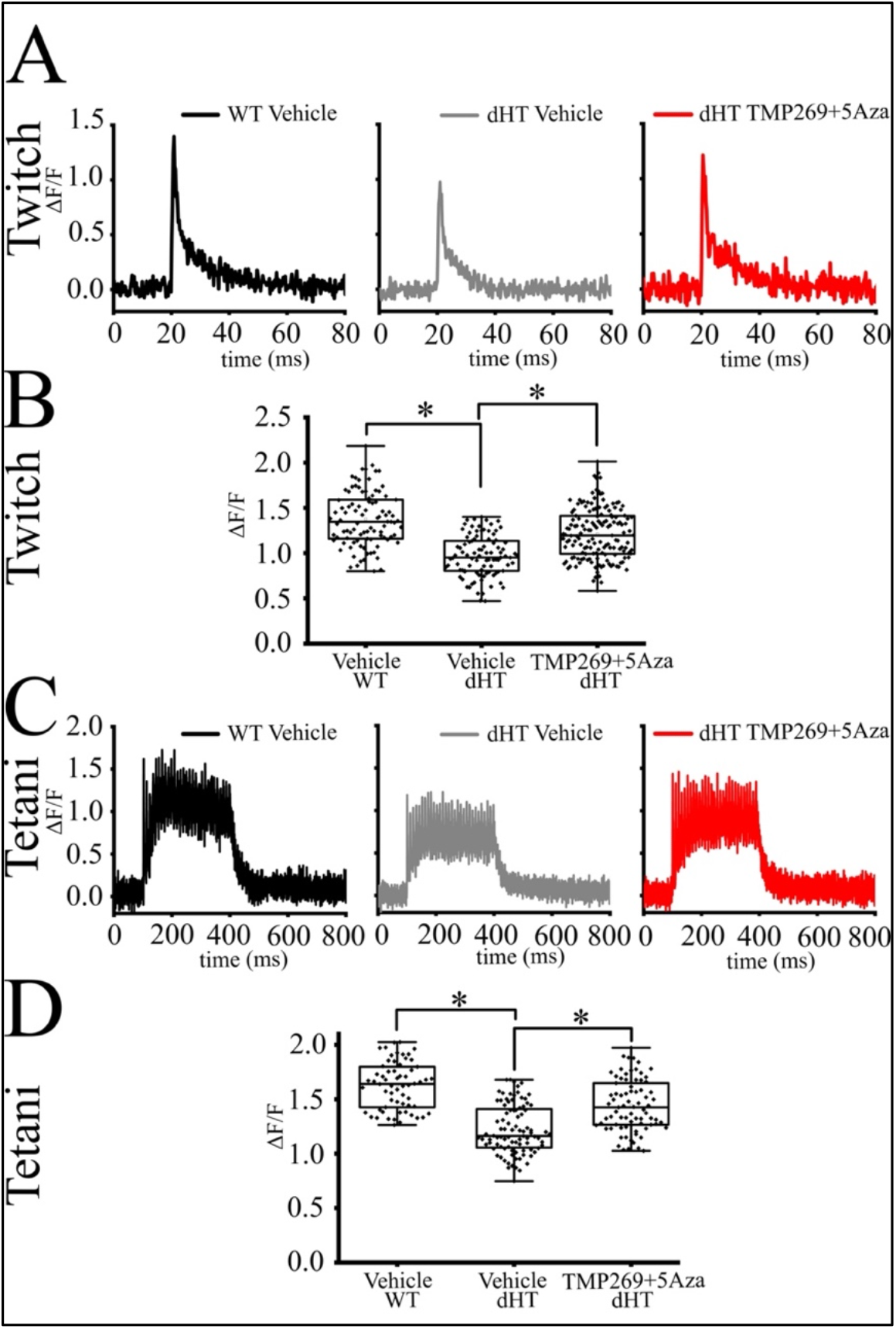
Electrically evoked peak Ca^2+^ transients in muscle fibers from treated dHT compound heterozygous (dHT) mice was rescued by TMP269+5Aza administration. Enzymatically dissociated FDB fibers dissected from 4-6 mice per group, were loaded with Mag-Fluo-4 and electrically stimulated by field stimulation. Black line, vehicle treated WT, grey line, vehicle treated dHT, red line, TMP269+5Aza treated dHT. **A.** Representative Ca^2+^ transient evoked by a single pulse (twitch) of 50 V with a duration of 1 msec. **B.** Whisker plots of peak twitch. Each symbol represents results obtained from a single FDB fiber. **C.** Representative Ca^2+^ transient evoked by tetanic stimulation by a train of pulses delivered at 100 Hz for 300 msec. **D**. Whisker plots of peak transient induced by tetanic stimulation. Each symbol represents results obtained from a single FDB fiber. *p<0.05 (ANOVA followed by the Bonferroni post hoc test). The exact P values are given in Table S4.

### TMP269+5-Aza rescues RyR1 expression in muscles from dHT mice

The results obtained so far indicate that treatment with TMP269+5-Aza exerts a beneficial effect preferentially on slow twitch muscles. Nevertheless, the genome wide effects linked to the combined inhibition of class II HDACs and DNMT may affect a number of processes underlying muscle strength, making it difficult if not impossible to dissect the exact mechanisms underlying the improvement of slow twitch muscle function observed in dHT mice. However, based on our previous results, we postulate that the improvement of muscle strength observed in soleus muscles may be explained, at least in part, by an increase of the key proteins involved in skeletal muscle activation. Fig. 5A shows Ryr1 transcript expression in soleus muscles from WT and dHT mice. Treatment with vehicle alone does not rescue Ryr1 expression (Mann-Whitney two-tailed test, WT vehicle vs dHT vehicle, P=0.039), however treatment with TMP269+5-Aza for 15 weeks causes a significant increase in Ryr1 transcript levels (Mann-Whitney two-tailed test, dHT vehicle vs dHT TMP269+5-Aza, P=0.019). Cacna1s levels are not affected by vehicle or drug treatment. Hdac4 transcript levels are increased in soleus muscles from dHT vehicle treated mice, compared to vehicle treated WT mice (Fig. 5A, Mann-Whitney two-tailed test, WT vehicle vs dHT vehicle, P=0.041). Hdac4 transcript levels decrease in muscles from TMP269+5-Aza treated dHT mice compared to vehicle treated dHT mice (Fig. 5A, Mann-Whitney two-tailed test, dHT vehicle vs dHT TMP269+5-Aza, P=0.002). We also investigated RyR1 protein content in total homogenates from soleus muscles from WT and dHT mice. The RyR1 protein content of soleus muscles from dHT mice injected with vehicle is 46% lower compared to that of WT mice (Fig. 5B). The mean ±S.D. % intensity of the immunopositive band corresponding to RyR1 is 100%±7.30, n=8 in WT vs 54.93%±18.45, n=6 in dHT, respectively (ANOVA followed by the Bonferroni post hoc test, *P=0.039). The RyR1 protein content is partially re-established by treatment with TMP269+5-Aza. Indeed, the RyR1 protein content in drug treated dHT mice increased and reaches a value of 88.03%±10.42*, n=7, in contrast to the 54.93%±18.45, n=6 found in vehicle treated dHT mice (ANOVA followed by the Bonferroni post hoc test, *P=0.037 for vehicle vs TMP269+5-Aza treated dHT mice; Fig. 5B). This recovery of RyR1 is consistent with the *in vivo* and *in vitro* muscle phenotype amelioration induced by the combined drug treatment. We also measured the effect of the combined drug treatment on the content of other proteins involved in skeletal muscle ECC including SERCA1, SERCA2, calsequestrin 1 and JP-45. As shown in Fig. 5C the muscle content of these proteins is unaffected by the drug treatment.

**Figure 5.**
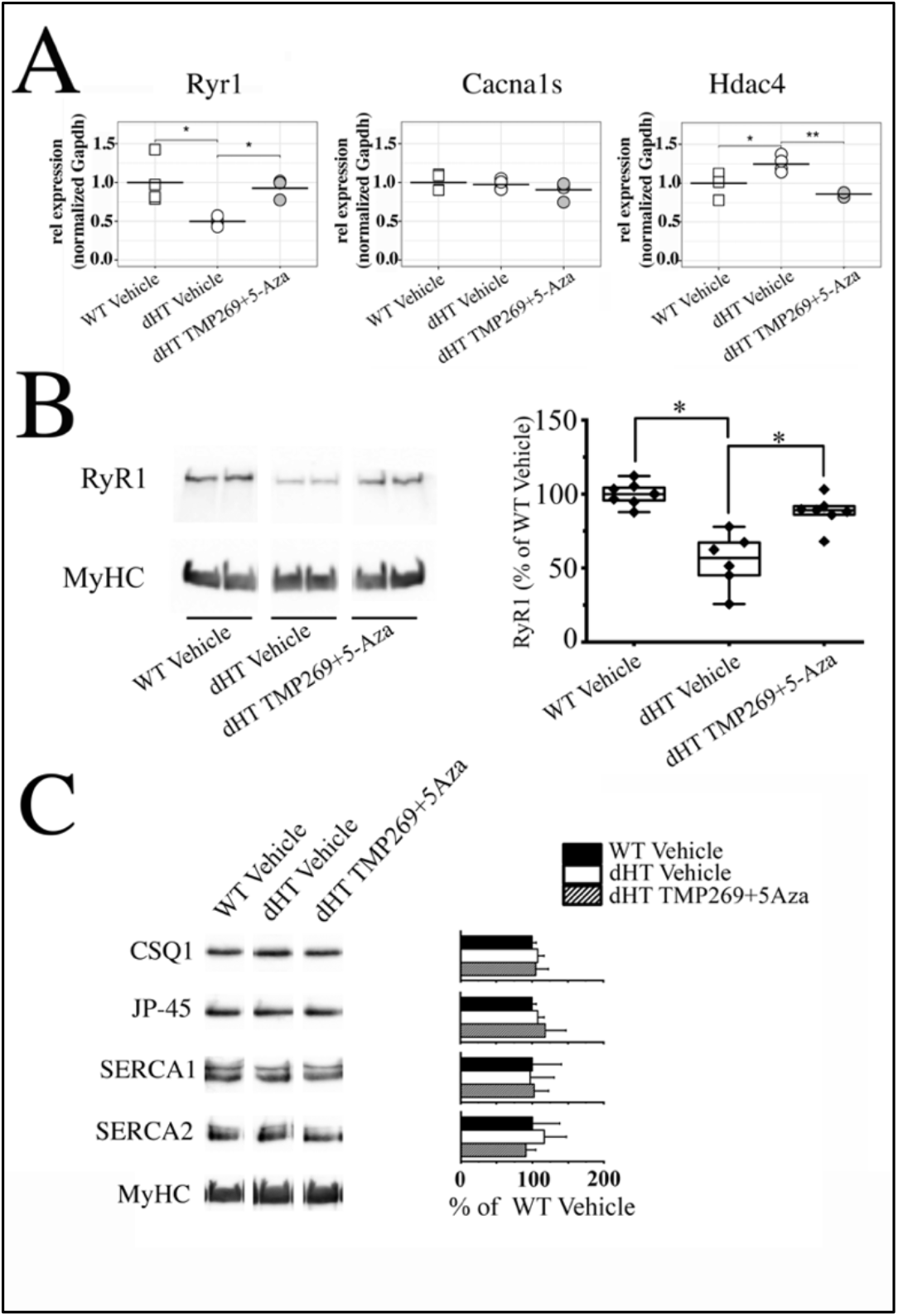
Treatment with TMP269+5Aza reverses RyR1 loss in soleus muscles from dHT mice. **A**. Real time qPCR on RNA isolated from soleus muscles isolated from vehicle treated WT and vehicle treated and TMP269+5-Aza treated dHT mice. Experiments were carried out on muscles isolated from 4 mice per group. RNA isolation and amplification conditions as described in the Methods section. *p<0.05; **p<0.01 ANOVA followed by the Bonferoni post hoc test. **B.** Western blot analysis of RyR1 content in total homogenates of soleus muscle from WT and dHT mice treated with vehicle or TMP269+5Aza. Proteins were separated on a 6 % SDS PAGE, blotted overnight onto nitrocellulose. Left panel: representative images of blot probed with anti-RyR1 Ab followed by anti-MyHC Ab for normalization. Right panel, data points are expressed as Whisker plots *p<0.05 (ANOVA followed by the Bonferroni post hoc test). **C** (left panel) representative immunoblots of total homogenates soleus muscles from WT (vehicle) and dHT (vehicle and TMP269+5Aza) mice probed with the indicated antibodies. (Right panel) bar histograms showing the mean ± S.D. (n=4 mice per group) intensity of the immunopositive band, expressed as % of the intensity of the band in WT (vehicle treated) mice. MyHC was used for loading protein normalization. Forty μg of protein were loaded per lane and proteins were separated on 7.5% or 10% SDS-PAG. Exact P values are given in the text.

### Recovery of calcium release units (CRUs) in soleus muscles from dHT after TMP269+5-Aza treatment

Skeletal muscle fibers from adult wild type (WT) are usually characterized by a regular transverse pale-dark striations. Within the fiber interior Ca release units (CRUs), the SR-TT junctions containing RyR1, are uniformly distributed, and mostly placed at A-I band transition (when sarcomeres are relaxed), on both sides of the Z line. CRUs are formed by two SR terminal cisternae closely opposed to a central TT oriented transversally with respect to the longitudinal axis of the myofibrils. These CRUs are called triads. CRUs are often associated with a mitochondrion, to form functional couples (24).

In soleus muscle fibers from vehicle treated dHT mice (Fig. 6A and B) CRUs present some abnormal features: the SR is often dilated (small arrows in Fig. 6A) and at times they are incomplete, meaning they are formed by only 2 elements (dyads) (Fig. 6B, black arrow). Quantitative EM analysis shows that in fibers of vehicle treated dHT mice there are 40.3±2.4 CRUs/100 μm^2^, 14.2±3.6% of them being dyads (Table 1, columns A-B). In soleus muscle fibers of dHT vehicle treated mice there are also regions with accumulated autophagic material (Fig. 6B, empty arrow). In fibers from TMP269+5-Aza treated dHT mice (Fig. 6C and D), analysis of CRUs indicates a (at least partial) rescue of the EC coupling machinery. Dilated SR in triads (pointed by small arrows in Fig. 6A) are practically absent (Fig. 6C)), and CRUs are more abundant and better preserved, meaning that they have the classic triad structure (3 elements: 2 SR and one T-tubule, small arrows in Fig. 6D). Quantitative analysis indicates that there are 46.9 ± 2.3 CRUs/100 μm^2^, only 6.5 ± 0.6% of them being dyads (Table 1, columns A-B).

**Figure 6.**
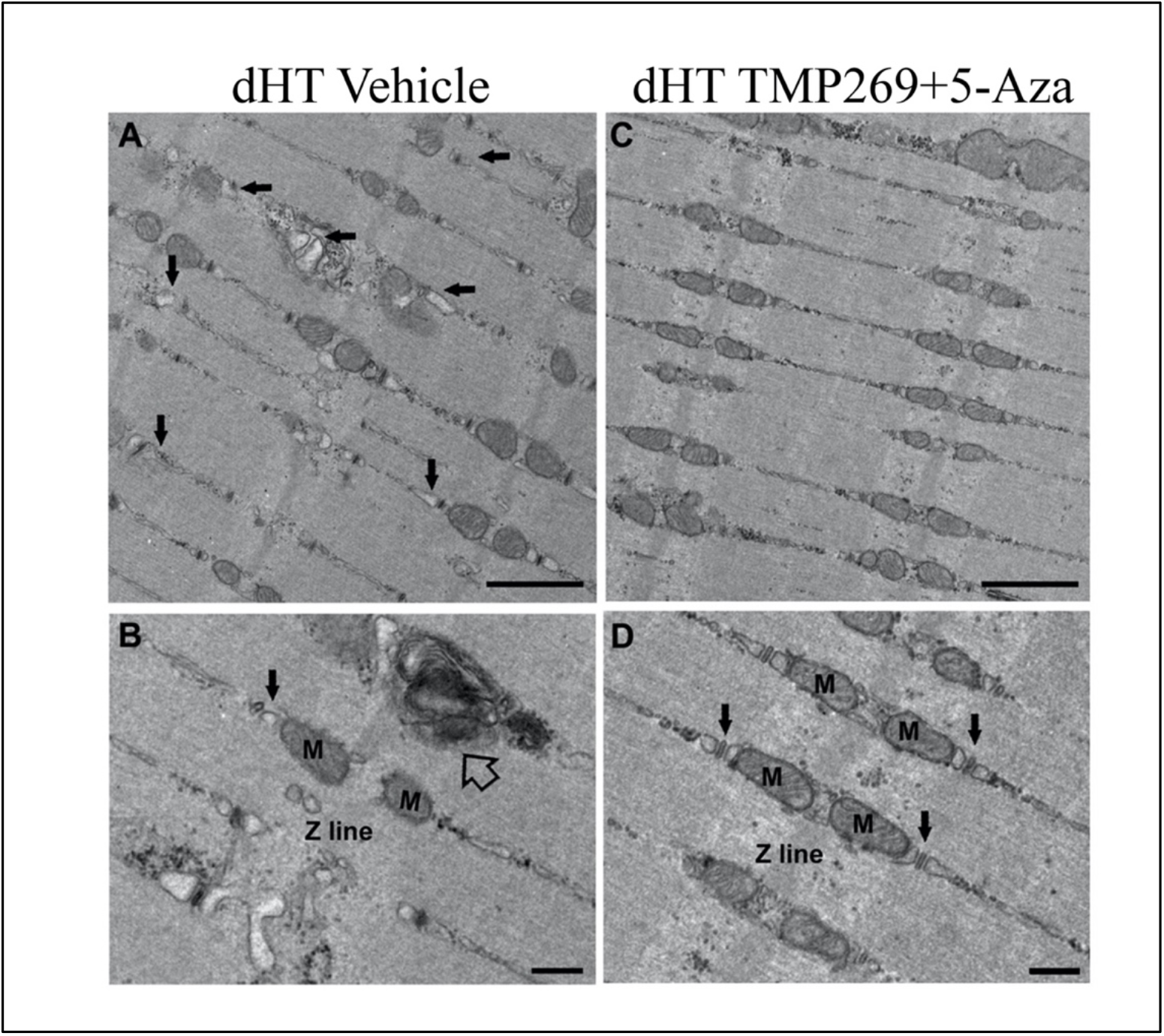
Representative EM images of soleus fibres from vehicle treated (A and B) and TMP269+5-Aza treated (C and D) dHT mice. **A.** Representative EM images at low magnification of soleus fibres from vehicle treated and **C.** TMP269+5Aza treaded dHT mice: small arrows point to dilated SR. **B.** and **D.** Higher magnification images showing calcium release units (CRUs, black arrow) and autophagic material (empty arrow, panel B) from vehicle treated and drug treated (D) dHT mice. M = mitochondria. Scale bars: A and C, 1 μm; B and D, 500 nm.

**Table 1.**
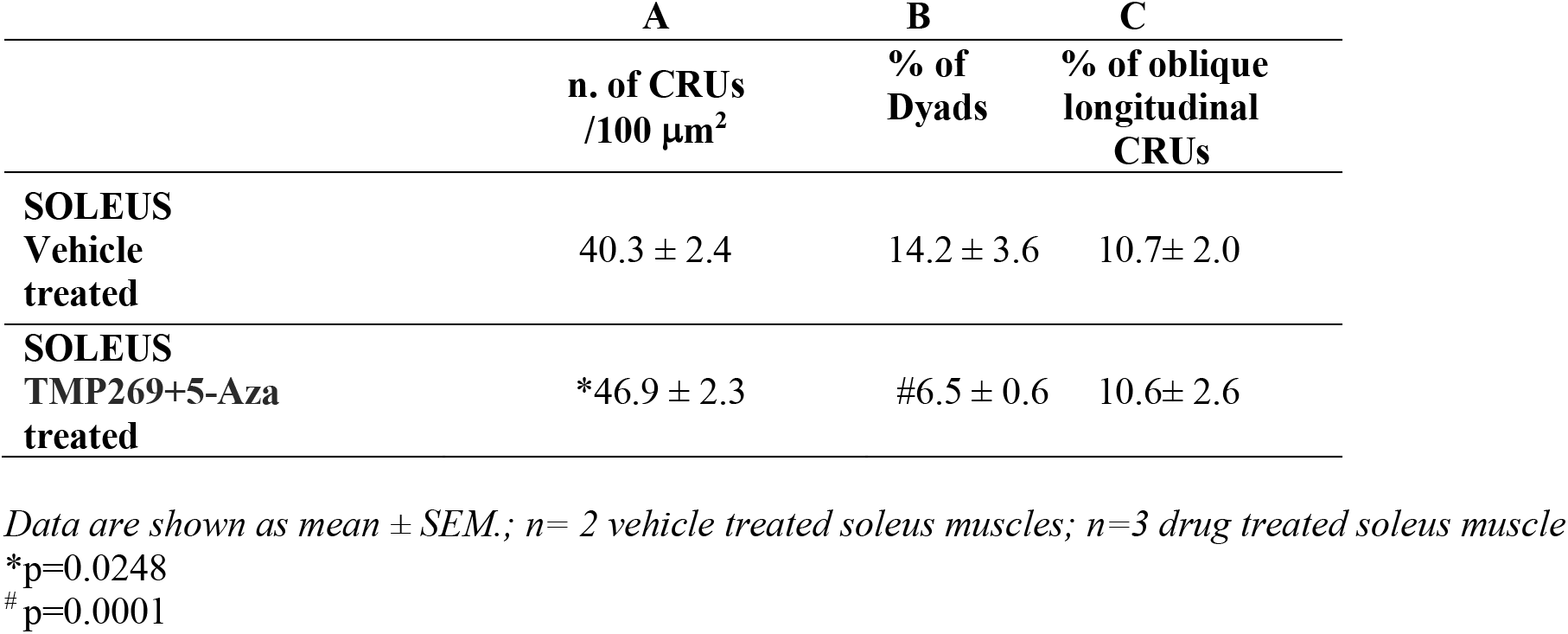
Quantitative analysis of CRUs. In soleus fibres from TMP269+5-Aza treated dHT mice, frequency of total CRUs and dyads (incomplete CRUs) are significantly rescued (columns A and B), suggesting a better preservation of CRUs and improved structure of triads.

We also analyzed mitochondria and their association with CRUs (Table 2). In TMP269+5-Aza treated dHT versus vehicle treated mice: i) the number/area of mitochondria is significantly increased (71.0±3.5 v.s 59.7±2.9 mean±S.E.M; Table 2, column A); ii) the number of damaged mitochondria is slightly but significantly reduced ( 2.9 ± 0.6 vs. 3.2 ± 0.5 mean±S.E.M; Table 2, column B), and finally iii) mitochondria are more often correctly associated with CRUs to form functional couples (34.5 ± 2.3 vs. 26.9 ± 2.2 mean±S.E.M; Table 2, column C).

**Table 2.** Quantitative analysis of mitochondria. Soleus fibres from TMP269+5-Aza treated dHT mice show a significant improvement in frequency, disposition and morphology of mitochondria.

|  | A | B | C |
| --- | --- | --- | --- |
| | n. of<br>mitochondria<br>in 100 $\mu\text{m}^2$ | n. of<br>severely altered<br>mitochondria in 100<br>$\mu\text{m}^2$ (%) | n. of<br>mitochondria<br>/CRU pairs<br>in 100 $\mu\text{m}^2$ |
| **SOLEUS**<br>**Vehicle Treated** | 59.7 ± 2.9 | 3.2 ± 0.5 (6.2) | 26.9 ± 2.2 |
| **SOLEUS**<br>**Drug Treated** | * 71.0 ± 3.5 | #2.9 ± 0.6 (3.7) | §34.5 ± 2.3 |

Data are shown as mean ± SEM.; n= 2 vehicle treated soleus muscles; n=3 drug treated soleus muscles.
\*p=0.0175
#p=0.0475
§p=0.0201

## Discussion

Congenital myopathies are rare neuromuscular disorders, caused in approximately 30% of the patients, by mutations in the *RYR1* gene (6, 8). Patients can present a variety of symptoms and phenotypes depending on whether the mutations are dominantly or recessively inherited. In particular, patients with recessive *RYR1* mutations often display involvement of extraocular muscles leading to ophthalmoplegia and or ptosis as well as involvement of respiratory muscles, often requiring assisted ventilation (6–8, 11, 15). Children may also present physical abnormalities including club foot, scoliosis, facial dysmorphisms, winged scapula and/or pectus excavatum (15). Nevertheless, the most disturbing symptom present at infancy in children carrying recessive *RYR1* mutations are hypotonia and proximal muscle weakness. In the most severe cases, muscle weakness also impairs masticatory muscles causing dysphagia, whereby the affected infants are fed via percutaneous endoscopy gastrostomy, a procedure which influences the quality of life of the affected individuals and his/her family members. To date there are no therapies available for congenital myopathies.

In order to target these unmet clinical needs we exploited the heterozygous mouse model we created carrying compound heterozygous RyR1 mutations (p.Q1970fsX16+p.A4329D), for a preclinical study aimed at developing a therapeutic strategy for patients with congenital myopathies. In particular, we tested small molecules targeting epigenetic modifying enzymes such as class II HDACs and DNMT. Here we show that the combined treatment with TMP269 and 5-Aza-2 deoxycytidine have strong ergogenic effects on muscle functions, tested both *in vivo* and *in vitro*. The improvement of muscle performance is consistent with the increase of both peak calcium transients and RyR1 protein content in total muscle homogenates and is apparently not associated with changes in the fiber type composition or changes of the minimal Feret’s diameter of muscle fibers. We are confident that this study provides proof of concept for a therapeutic strategy aimed to enhance muscle strength in patients affected by congenital myopathies linked to recessive *RYR1* mutations.

Our data shows that the improvement of muscle performance *in vivo* is mainly due to the amelioration of muscle strength in slow twitch fibers. The muscle ultrastructure demonstrates a partial rescue of the EC coupling machinery (more triads, less of them being incomplete) and of the associated mitochondria and may explain in part the rescue of muscle function. The elucidation of the exact molecular mechanisms underlying the improvement in strength specifically of slow twitch fibers caused by genome wide effects of drugs inhibiting epigenetic modifying enzymes is an arduous challenge. However, we believe that a few assumptions can be taken into account to understand our data. First, histopathology of muscle biopsies from patients harboring recessive *RYR1* mutations shows fiber type I predominance (15); such an observation suggests that recessive *RYR1* mutations have a greater damaging effect on fast twitch fibers compared to slow twitch fibers. This interpretation of the histopathology of human skeletal muscle biopsies is consistent with our previous results on the dHT mouse model (20). Indeed, we found that the RyR1 array disarrangements and morphological alterations are more noticeable in fast glycolytic muscles such as EDLs compared to FDB and soleus muscles, a result in line with the idea that skeletal muscle oxidative fibers are more resistant to the damaging effects linked to the expression of recessive *RYR1* mutations (20). If this is the case, then upon pharmacological treatment slow oxidative fibers should recover their function to a greater extent than fast twitch glycolytic fibers. Second, inhibition of class II HDAC activity by genetic manipulation leading to the inactivation of all class II HDAC genes promotes skeletal muscle endurance capacity by activating Mef2 dependent gene transcription (25). We found that HDAC4 transcript levels in soleus muscles from dHT mice treated with a combination of TMP269+5-Aza decreases to a value similar to that found in WT littermates. Though we don’t have a clear explanation for this observation, it appears to be in line with the proposal that high levels of class II HDACs are indicative of muscle damage or age-related atrophy and reduced expression of class II HDACs is associated with an increased running endurance (25, 26). Third, it was recently demonstrated that pharmacological inhibition of class II HDACs causes de-acetylation not only of nuclear histones but also of other targets, including MyHC and PGC1alfa (27), an event which causes an increase of the cytoplasmic level of the PGC1alpha protein. Interestingly, PGC1alpha is a crucial regulator of oxidative metabolism and mitochondrial biogenesis in skeletal muscle fibers (28). We found that administration of class II HDAC and DNMT inhibitors to dHT mice induces an increase in the number of mitochondria in slow twitch fibers. This result is in agreement with the observed increase in PGC1alpha activity observed after treatment with class II specific HDAC inhibitors (27). Fourth, reconstitution of the RyR1 arrays in the junctional SR might restore the RyR1 retrograde signal, which is important for the activation of Cav1.1 Ca^2+^ currents across T tubular membranes. Since the Cav1.1 to RyR1 ratio is lower in slow twitch muscles (29), reactivation of the Cav1.1 calcium current might occur to a larger extent in slow twitch muscles from dHT treated mice compared to fast twitch muscles. Though Cav1.1 calcium currents are not relevant for skeletal muscle EC coupling, they have been implicated in a broad range of muscle functions, including fatigue resistance, protein synthesis and calcium dependent signaling pathways involved in the maintenance of proper skeletal muscle function (30).

In conclusion, the present study demonstrates that the combined pharmacological treatment with an FDA approved DNMT inhibitor and TMP269 improves muscle strength and performance in a mouse model for recessive *RYR1* congenital myopathy. Further studies are needed to define the exact mechanism of the ergogenic effects of these genome wide epigenetic modifying drugs; nevertheless, our results provide the proof of concept for the development for pharmacological treatment of patients with congenital myopathies linked to recessive *RYR1* mutations.

## Materials and Methods

### Compliance with Ethical standards

All experiments involving animals were carried out on 16-21 weeks old male mice unless otherwise stated. Experimental procedures were approved by the Cantonal Veterinary Authority of Basel Stadt (BS Kantonales Veterinäramt Permit numbers 1728 and 2950). All experiments were performed in accordance with ARRIVE guidelines and Basel Stadt Cantonal regulations.

### Drug injection protocol

The class II HDAC inhibitor TMP269 was purchased from Selleckchem (S7324), the DNA methyltransferase inhibitor 5-aza-2 deoxycytidine was from Sigma-Aldrich (A3656), Polyethylenglycol 300 (PEG300) was from Merck (8.17019), N-Methyl-2-pyrrolidone (NMP) was from Sigma-Aldrich (328634). Intraperitoneal injections (IP, 30-gauge needle) started at the sixth week of age and continued daily for 10 - 15 weeks. Mice received vehicle (PEG300 and NMP), TMP269 (25 mg/Kg), 5-aza-2-deoxycytidine (0.05 mg/Kg), a combination of the two compounds (TMP26+5-aza-2-deoxycytidine, 25 mg/Kg and 0.05 mg/Kg, respectively). TMP269 and 5 aza-2-deoxycytidine were diluted in PEG300 (500 μl/Kg) and NMP (250 μl/Kg). The final volume of drug or vehicle injected per mouse was 750 μl per kilogram body weight.

### Pharmacokinetics analysis of TMP269

Six weeks old mice were injected intraperitoneally with 25 mg/kg bodyweight (n=4) TMP269 and blood and muscle tissue were collected at different time points post injection and analyzed. Blood (20 μL) was collected from the tail vein using lithium heparin coated capillaries (Minivette POCT 20 μL LH, Sarstedt, Germany). Samples were collected prior treatment (T0) and 1, 3, 6, 12, 24, 48 hours after TMP269 injection, transferred directly into autosampler tubes, and kept at −20°C until analyzed. For skeletal muscle samples, the following protocol was used. At T=0 and at different time points after TMP269 injection (1, 3, 12, 24, 48 hours), mice were sacrificed and their skeletal muscles were isolated, flash frozen and stored in liquid nitrogen. On the day of the analysis, muscle samples were homogenized using a tissue homogenizer (Mikro-Dismembrator S, Sartorius, Aubagne, France) for two sequences of 30 sec each.

The levels of TMP269 in blood and in skeletal muscle were quantified by high pressure liquid chromatography tandem mass spectrometry (LC-MS/MS) as described (21). Briefly, A Shimadzu LC system (Kyoto, Japan) was used, TMP269 and TMP195 (internal standard, IS) were analyzed by positive electrospray ionization and multiple reaction monitoring (MRM). Methanol containing IS (5 nM TMP195) was used to extract TMP269 from blood and muscle homogenate samples (20 μL blood/muscle plus 175 μL methanol). The samples were vortex mixed for approximately 30 seconds and centrifuged at 10°C, 3220 g for 30 minutes. Ten μL of supernatant were injected into the LC-MS/MS system. Calibration lines were prepared in blank mouse blood and covered a range of concentrations, from 0.25 nM to 500 nM TMP269. A linear regression between TMP269 to the IS peak area ratio and the nominal concentration was established with a weighting of 1/x2 to quantify the TMP269 concentration in blood or muscle. The Analyst 1.6.2 software (AB Sciex, Concord, Canada) was used to operate the LC-MS/MS system and to analyze the data.

### Genotyping dHT mice and real time qPCR

Mouse genotyping was performed on mouse genomic DNA, by PCR amplification of *Ryr1* exon 36 and exon 91 using specifically designed primers (Supplementary Table S5), as previously described (20). Quantitative real-time polymerase chain reaction (qPCR) was carried out on RNA extracted from muscles isolated from 21 weeks old mice (treated with drug or vehicle for 15 weeks, starting at 6 weeks of age). Briefly, total RNA extracted from frozen hind limb muscles was isolated, genomic DNA was removed by DNase I treatment (Invitrogen; 18068-015) and 1000 ng RNA were reverse-transcribed into cDNA using the High capacity cDNA Reverse Transcription Kit (Applied Biosystems; 4368814) as previously described (20). The cDNA was amplified by qPCR using the primers listed in Supplementary Table S5 and transcript levels were quantified using Power SYBR^®^ Green reagent Master Mix (Applied Biosystems; 4367659), using the Applied Biosystems 7500 Fast Real-time PCR System running 7500 software version 2.3 as previously described (20). Transcript quantification was based on the comparative ΔΔCt method. Each reaction was performed in duplicate and averaged and results are expressed as relative gene expression normalized to the housekeeping gene glyceraldehyde 3-phosphate dehydrogenase (Gapdh).

### *In Vivo* Muscle Strength Assessment

*In vivo* muscle performance was evaluated in male mice, by performing the following measurements: (i) forelimb grip strength and (ii) spontaneous locomotor activity. Forelimb grip strength was assessed once per week for a period of 10 weeks using a Grip Strength Meter from Columbus Instruments (Columbus, OH, USA), following the manufacturer’s recommendations. The grip force value obtained per mouse was calculated by averaging the average value of 5 measurements obtained on the same day on the same mouse. To avoid experimental bias during measurements of grip force, experimenters were blinded to the genotype and treatment of mice. For spontaneous locomotor activity, after 15 weeks of treatment with vehicle or TMP269+ 5-Aza, mice were individually housed in cages equipped with a running wheel carrying a magnet as previously described (20, 31). Wheel revolutions were registered by reed sensors connected to an I-7053D Digital-Input module (Spectra), and the revolution counters were read by a standard laptop computer via an I-7520 RS-485-to-RS-232 interface converter (Spectra). Digitized signals were processed by the ‘‘mouse running’’ software developed at Santhera Pharmaceuticals (32). Total running distance (kilometer) and speed (Km/h) were evaluated.

### *Ex vivo* Muscle Strength Assessment

To test muscle force ex vivo, *extensor digitorum longus* (EDL) and soleus muscles were dissected from 21 weeks old male WT and dHT mice, after 15 weeks of treatment with vehicle alone, or with the combination of TMP26915-Aza. Isolated EDL and soleus muscles were mounted onto a muscle force transducing setup (MyoTronic Heidelberg) as previously described (31). Muscle force was digitized at 4 kHz by using an AD Instrument converter and stimulated with 15 V pulses for 1.0 msec. Tetanus was recorded in response to a train of pulses of 400 msec and 1100 msec duration delivered at 10/20/50/100/150 Hz, and 10/20/50/100/120 Hz, for EDL and soleus, respectively. Specific force was normalized to the muscle cross-sectional area [CSA_wet weight (mg)/length (mm)_1.06 (density mg/mm^3^)] (33). The experimenter performing the measurements was blinded with respect to the mouse genotype and treatment.

### Isolation of single *flexor digitorum brevis* (FDB) fibers for intracellular Ca^2+^ measurements

Twenty-one weeks old WT and dHT vehicle and drug treated male mice were killed by pentobarbital overdose according to the procedures approved by the Kantonal Veterinary Authority. *Flexor digitorum brevis* (FDB) muscles were isolated and digested with 0.2% of Collagenase type I (Clostridium hystoliticum Type I, Sigma-Aldrich) and 0.2% of Collagenase type II (Clostridium hystoliticum Type II, Worthington) in Tyrode’s buffer (137 mM NaCl, 5.4 mM KCl, 0.5 mM MgCl_2_, 1.8 mM CaCl_2_, 0.1% glucose, 11.8 mM HEPES, pH 7.4 NaOH) for 45 minutes at 37 °C as described (34). Muscles were washed with Tyrode’s buffer to block the collagenase activity and gently separated from tendons using large to narrowest set of fire-polished Pasteur pipettes. Fibers obtained by this procedure remained excitable and contracted briskly when assayed experimentally. Finally, fibers were placed on laminin coated (5 μl of 1 mg/ml mouse laminin from ThermoFischer) 35 mm glass bottom dishes (MatTek corporation) for measurements of the resting [Ca^2+^] or on ibiTreat 15μ-Slide 4 well (Ibidi) for electrically evoked Ca^2+^ measurements as previously described (20).

### Measurements of resting [Ca^2+^] and of electrically evoked Ca^2+^ transients

Single FDB fibers were isolated from 4-5 male mice per group and allowed to adhere to laminin treated 35 mm glass bottom dishes for 1 hr at 37°C. The fibers were then loaded with 5 μM Fura-2 AM (Invitrogen) by incubating them for 20 min at 19°C in Tyrode’s buffer. The excess Fura-2 was diluted out by the addition of fresh Ringer’s solution and measurements of the resting [Ca^2+^] were carried out using an inverted Zeiss Axiovert fluorescent microscope, as previously described (20, 31). Only those fibers that contracted when an electrical stimulus was applied were used for the [Ca^2+^] measurements.

For electrically evoked Ca^2+^ transients, single FDB fibers were incubated for 10 min at 19 °C in Tyrode’s solution containing 10 μM low affinity calcium indicator Mag-Fluo-4 AM (Thermo Fischer), 50 μM N-benzyl-p-toluene sulfonamide (BTS, Tocris). Fibers were rinsed twice with fresh Tyrode’s solution, and measurements were carried out in Tyrode’s solution containing 50 μM BTS. Measurements were carried out with a Nikon Eclipse inverted fluorescent microscope equipped with a 20x PH1 DL magnification objective. The light signals originating from a spot of 1 mm diameter of the magnified image of FDB fibers were converted into electrical signals by a photomultiplier (Myotronic Heidelberg). Fibers were excited at 480 nm and then stimulated either with a single pulse of 50 V with a duration of 1 msec, or with a train of pulses of 50 V with a duration of 300 msec delivered at 100 Hz. Fluorescent signals were acquired using PowerLab Chart7. Changes in fluorescence were calculated as ΔF/F0= (Fmax-Frest)/(Frest). Kinetic parameters were analyzed using Chart7 software. FDB fibers were isolated from 4-6 mice per group and results were averaged.

### Biochemical analysis of total muscle homogenates

Total muscle homogenates were prepared from soleus muscles isolated from WT and dHT vehicle and drug treated mice. SDS-polyacrylamide electrophoresis and Western blots of total homogenates were carried out as previously described (20, 31, 35). Western blots were stained with the primary antibodies listed in Supplementary Table S6, followed by peroxidase-conjugated Protein G (Sigma P8170; 1:130’000) or peroxidase-conjugated anti-mouse IgG (Fab Specific) Ab (Sigma A2304; 1:200’000). The immuno-positive bands were visualized by chemiluminescence using the WesternBright ECL-HRP Substrate (Witec AG). Densitometry of the immune-positive bands was carried out using the Fusion Solo S (Witec AG). A representative immunoblot of each antibody on total muscle homogenates from WT mice is shown in Supplementary Figure S3.

### Genome-wide DNA Methylation Analysis

Sodium-bisulfite treated DNA of nine samples (6 dHT vehicle treated and 3 dHT drug treated) was subjected to measure global DNAm by *Infinium Mouse methylation* BeadChip array (Illumina) according to manufacturer’s instructions. Illumina mouse array contains ~285,000 methylation sites per sample at single-nucleotide reslution covering over 20 design categories including gene promoters and enhancers. Downstream analysis were conducted using annotation developed previously mapping to mm10. Raw IDAT files were preprocessed using R/Bioconductor package SeSAMe (36). Samples were normalized using “noob” including background subtraction and dye-bias normalization. Methylation levels were computed as Beta (β) values (Meth / [Meth + Unmeth +100]) and as logit-transformed M-value (log2 [Meth / Unmeth]). Probes where at least one sample had a detection *P* value > 0.05 and the probes recommended previously using mm10 genome as general-purpose masking probes were filtered out (36). The remaining 215,717 CpGs were used for further analysis. Statistical analyses were performed on M-values (37) using bioconductor *limma*, whereas β-values were used for biologic interpretation. Normalized methylation levels were evaluated using linear regression model between dHT treatment vs. vehicle. Differentially methylated CpGs were identified using empirical Bayes moderated t-statistics and associated Benjamini-Hochberg (BH) false discovery rate (FDR) adjusted *P*-values <0.05 were used as cutoff.

### Histological examination

Soleus muscles from treated and untreated WT and dHT mice were isolated and mounted for fluorescence microscopy imaging. Muscles were embedded in OCT and deep-frozen in 2-Methylbutane, then stored at minus 80°C. Subsequently, transversal 10 μm sections were obtained using a Leica Cryostat (CM1950) starting from the belly of the muscles. Sections were stained as described by Delenzie et al. 2019 (38) using the following primary antibodies (see Supplementary Table S5 for suppliers and catalog numbers): mouse IgG2b anti-MyHC I (1:50), mouse IgG1 anti-MyHC IIa (1:200), mouse IgM anti-MyHC IIb (1:100) and rabbit anti-mouse laminin (1:1500), followed by incubation with the following secondary antibodies: goat anti-mouse IgG1Alexa Fluor 568 (1:1000), goat anti-mouse IgM Alexa Fluor 488 (1:1000), goat anti-mouse IgG Fcγ2b Dylight 405 (1:400) and Donkey anti-rabbit IgG Alexa Fluor 647 (1:2000). Images were obtained using an Eclipse Ti2 Nikon Fluorescence microscope with 10X air objective lens. Muscles from 3 mice per group were evaluated. Images were analyzed using Fiji plugins developed by Delenzie et al. (38) in order to obtain information on fiber types and minimal Feret’s diameter (32), the closest possible distance between the two parallel tangents of an object, using a combination of cell segmentation and intensity thresholds as described (32).

### Preparation and quantitative analysis of samples by electron microscopy (EM)

Soleus muscles were dissected from sacrificed animals, pinned on a Sylgard dish, fixed at room temperature with 3.5% glutaraldehyde in 0.1 M NaCaCO buffer (pH 7.4), and stored in the fixative at 4°C (39). Fixed muscle were then post-fixed in a mixture of 2% OsO_4_ and 0.8% K_3_Fe(CN)_6_ for 1-2 h, rinsed with 0.1M sodium cacodylate buffer with 75 mM CaCl_2_, en-block stained with saturated uranyl acetate replacement, and embedded for EM in epoxy resin (Epon 812). Ultrathin sections (~40 nm) were cut in a Leica Ultracut R microtome (Leica Microsystem, Austria) using a Diatome diamond knife (DiatomeLtd. CH-2501 Biel, Switzerland) and examined at 60 kV after double-staining with uranyl acetate replacement and lead citrate, with a FP 505 Morgagni Series 268D electron microscope (FEI Company, Brno, Czech Republic), equipped with Megaview III digital camera (Munster, Germany) and Soft Imaging System (Germany).

### Quantitative analyses by EM

Data contained in Tables 1 and 2 were collected on soleus muscles from 21 weeks old dHT mice, either vehicle or TMP269+5-Aza treated. In each sample, 10-20 fibers were analyzed. In each fiber 2-3 micrographs (all at the same magnification, 14K, and of non-overlapping regions) were randomly collected from longitudinal sections.

Number of CRUs (Table 1, column A), number of mitochondria, number of severely altered mitochondria, and number of mitochondrion-CRU pairs (Table 2, columns A-C, respectively) were reported as average number/100 μm^2^ (24). In each EM image, we also determined the number of dyads, i.e. incomplete triads (Table 1, column B), and of oblique/longitudinal CRUs (Table 1, column C), and expressed as percentages over the total number of CRUs. Mean and SEM were determined using GraphPad Prism (GraphPad Software, San Diego, California USA). Statistically significant differences between groups were determined by the Student’s *t* test (GraphPad Software, San Diego, California USA) or by a Chi-squared test (GraphPad Software, San Diego, California USA). Values of p < 0.05 were considered significant.

### Statistical analysis

Statistical analysis was performed using the Student’s unpaired *t* test for normally distributed values when two groups were being compared, or the ANOVA test followed by the Bonferroni post hoc test for multiple group comparisons. The Mann–Whitney U test was used when values were not normally distributed. p<0.05 was considered significant.

## Supplementary Materials

**Fig. S1**. Pharmacokinetic profile of TMP269 following intraperitoneal injection in WT mice.

**Fig. S2**. Daily i.p injections with TMP269+5-Aza increase acetylation of Lys residues and of H3K9 in muscles from dHT mice.

**Fig. S3**. Immunoreactivity of the antibodies used in the study.

**Table S1**. List of hypomethylated protein-encoding genes in soleus muscles from dHT mice treated for 16 weeks with TMP269+5-Aza drug versus vehicle treated dHT mice.

**Table S2**. Specific force of EDL and soleus muscle from WT and dHT mice treated with vehicle or TMP269+5-Aza for 16 weeks.

**Table S3**. Fiber type composition of soleus muscles from mice treated for 16 weeks with vehicle or TMP269+5-Aza.

**Table S4**. Analysis of electrically evoked calcium transients in single FDB muscle fibers isolated from WT and dHT littermates, treated with vehicle or TMP269+5-Aza (25 mg/Kg) for 16 weeks

**Table S5**. Sequence of primers used and targets.

**Table S6.** List of antibodies and suppliers.

## List of Abbreviations

5-Aza: 5-Aza-2-deoxycytidine
CCD: central core disease
CRU: calcium release units
DHPR: dihydropyridine receptor
DNMT: DNA methyltransferase
ECC: excitation-contraction coupling
EDL: *extensor digitorum longus*
FDB: *flexor digitorum brevis*
HDAC: histone de-acetylase
MyHC: myosin heavy chain
MmD: multiminicore disease
SR: sarcoplasmic reticulum
TT: transverse tubules

## Author contributions

Conceptualization: FZ and ST

Methodology: AR, CB, ST, SB, FZ, MF, DU

Investigation: AR, ST, SB, MF, LP, SB, CB, FN, FP, FZ

Funding acquisition: ST, FZ

Project administration: ST, FZ

Supervision: ST, FZ, AR, FP

Writing – original draft: FZ, AR, ST

Writing – review & editing: FZ, ST, AR, SB, MF, LP, FZ, FP

## Data and materials availability

All data, code, and materials used in the analysis are available in some form to any researcher for purposes of reproducing or extending the analysis. There are no restrictions on materials, such as materials transfer agreements (MTAs). All data are available in the main text or the supplementary materials.

