## supplementary figures and tables for "Improvement of muscle strength in a mouse model for congenital myopathy treated with HDAC and DNA methyltransferase inhibitors"

**Supplementary files Ruiz et al.**

**Figure S1**

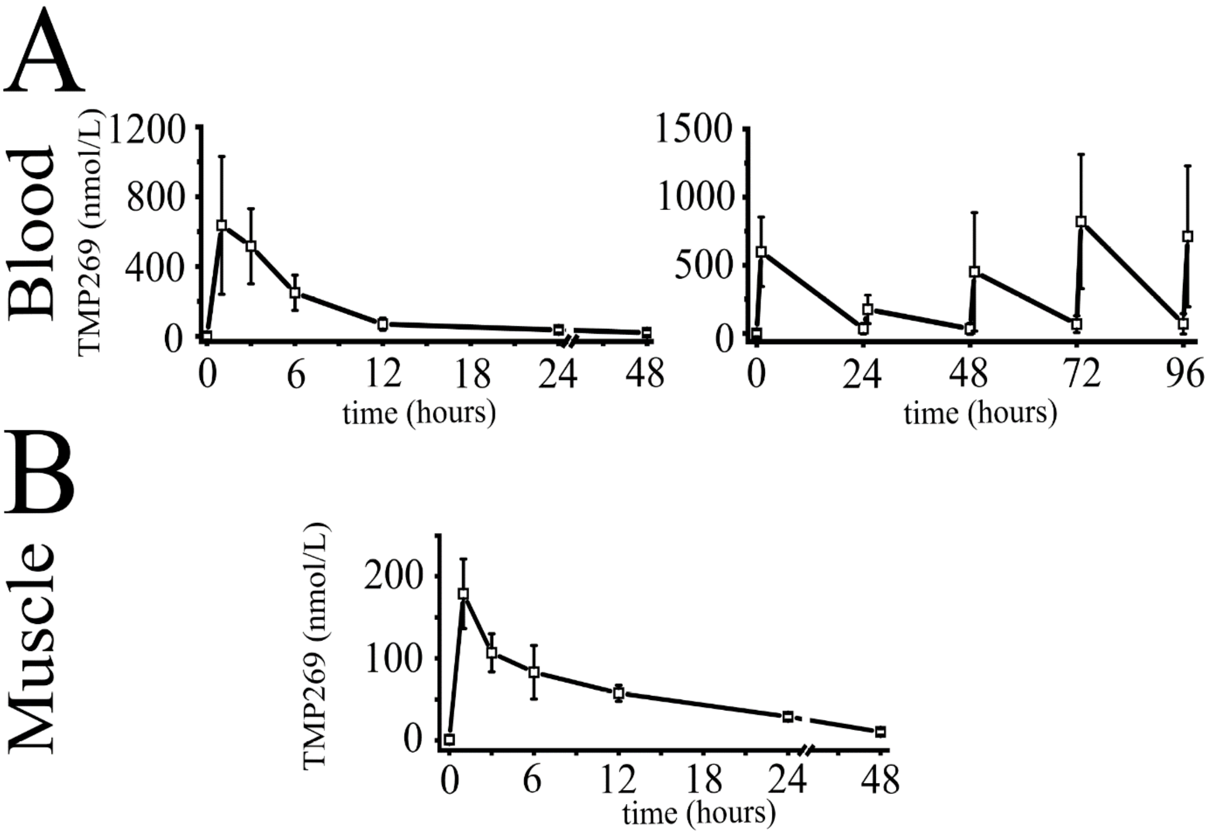


<u>Supplementary Figure S1</u>**: Pharmacokinetic profile of TMP269 following intraperitoneal injection in WT mice. A.** Concentration-time course of TMP269 in blood. Left panel: mean (±S.D., n=4 mice) plasma concentration in mice receiving a single dose of 25 mg/Kg of TMP269. Right panel, mean (±S.D., n=4 mice) blood concentration of TMP269 in mice after receiving consecutive doses of 25 mg/Kg of TMP269 at t=0, 24, 48, 72, and 96 hours. Blood samples were taken 10 min before and 1 hour after each intraperitoneal injection. **B.** Mean (±S.D.; n=4) TMP269 concentration in skeletal muscle after a single intraperitoneal injection of 25 mg/Kg TMP269, during 48 hours.

**Figure S2**

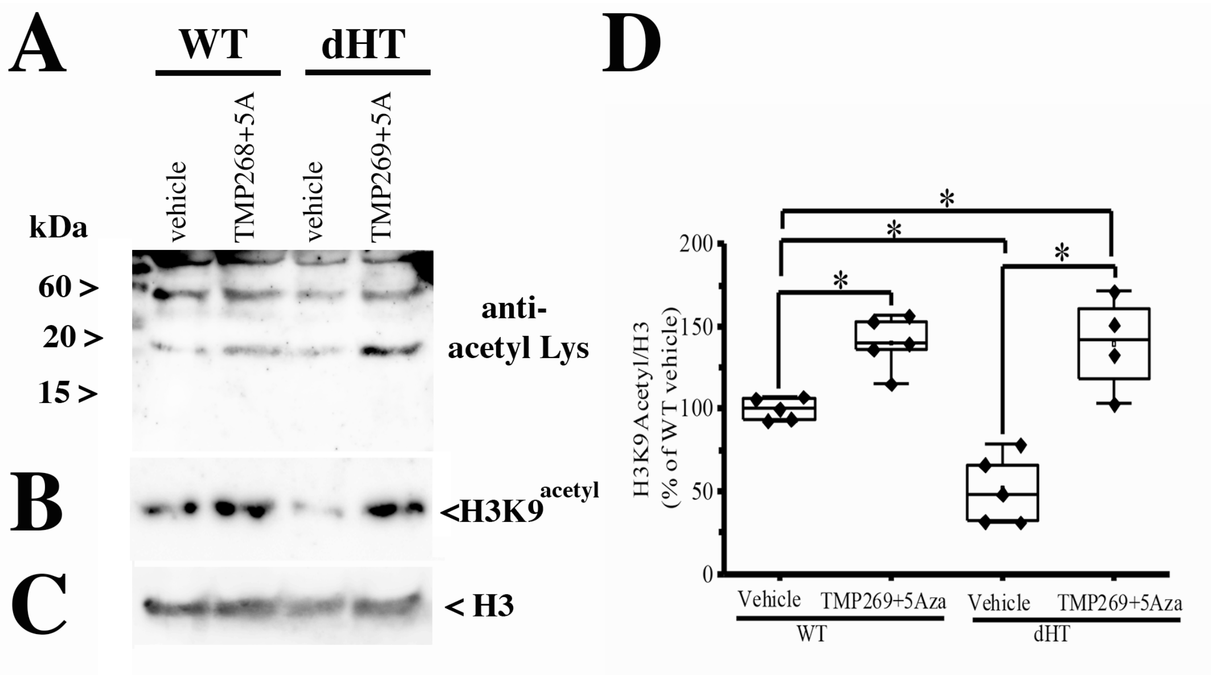


<u>Supplementary Figure S2:</u> **Daily i.p injections with TMP269+5-Aza increase acetylation of Lys residues and of H3K9 in muscles from dHT mice.** WT and dHT mice received a daily injection vehicle or 25 mg/kg TMP269+0.05 mg/Kg 5-Aza. After 15 weeks of treatment, approximately 200 FDBs fibers per mouse were isolated and resuspended in cracking buffer (10% glycerol, 5% ß-mercaptoethanol, 2.5% SDS, 62.5 mM Tris pH 6.8, 6 M Urea). Protein extracts were loaded on a Tris-Tricine gel, blotted onto nitrocellulose and probed with **(A)** anti-acetyl-Lys and (**B**) anti H3K9$^{acetyl}$ antibodies. **(C)** Shows the immunoreactivity of anti- H3 (histone 3) for loading control. **(D)** Boxplot analysis of western blots using anti-H3K9$^{acetyl}$ antibodies. The immunoreactivity of the H3K9$^{acetyl}$ positive band obtained in muscles from vehicle-treated WT mice was set to 100%. Each symbol represents the result obtained from a single mouse. *$p<0.05$ (ANOVA followed by the Bonferroni post hoc test).

WT vehicle vs WT TMP269+5-Aza, P= 0.041; WT vehicle vs dHT vehicle, P=0.037

WT vehicle vs dHT TMP269+5-Aza, P=0.042; dHT vehicle vs dHT TMP269+5-Aza, P=0.021

**Figure S3**

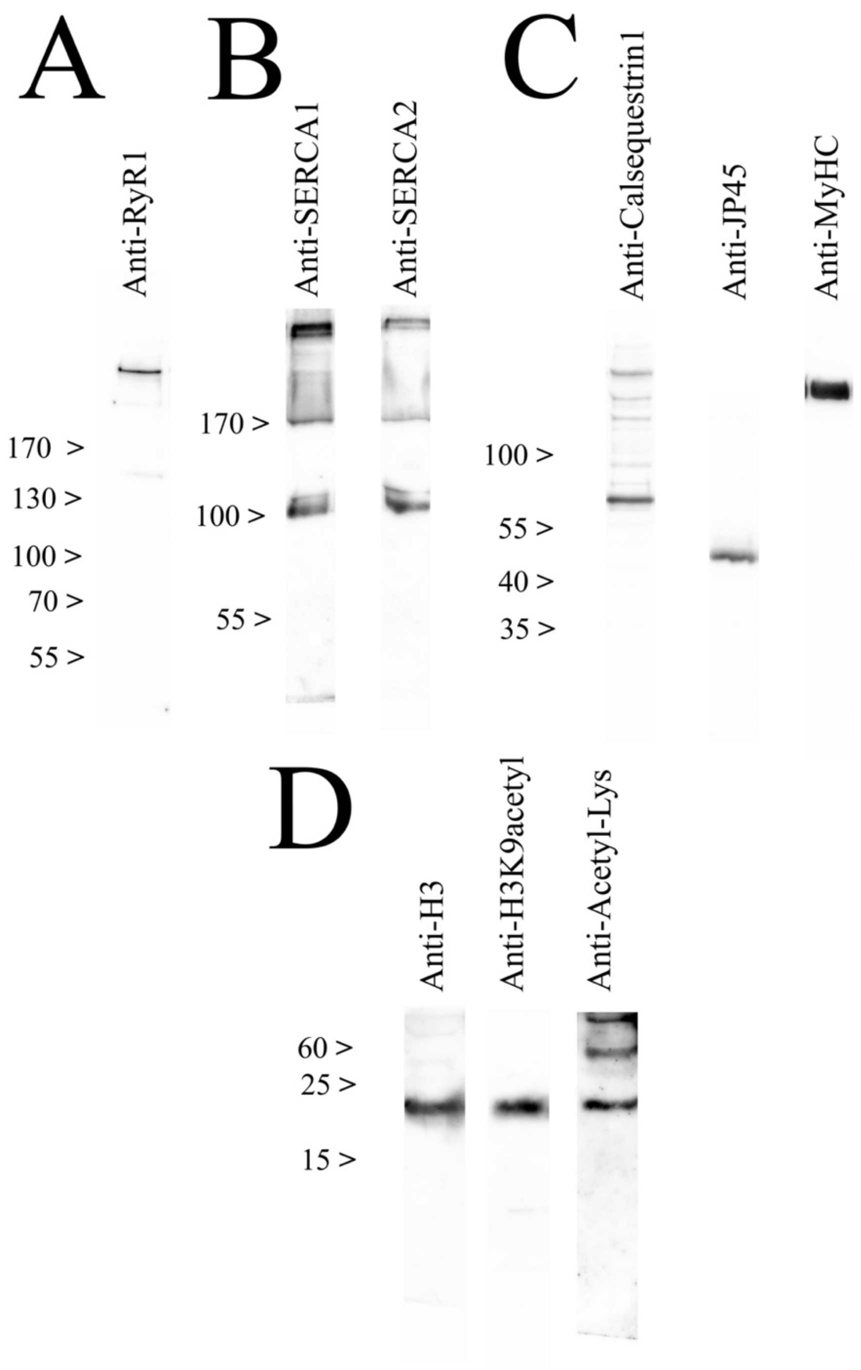


<u>Supplementary Figure S1</u>: **Immunoreactivity of the antibodies used in the study.** Total muscle homogenates were prepared from WT mice, proteins were separated on **(A)** 6 % SDS PAG, **(B)** 7% SDS PAG **(C)** 10% SDS PAG and **(D)** Tris-Tricine gel 10%, and probed with the indicated primary antibodies. Blots were then incubated with peroxidase conjugated anti-mouse IgG or peroxidase conjugated protein-G and the immunopositive bands were visualized by chemiluminescence.

**Supplementary Table S1**. List of hypomethylated protein-encoding genes in soleus muscles from dHT mice treated for 15 weeks with TMP269+5-Aza drug versus vehicle treated dHT mice.

| Gene name | Gene name | Gene name | Gene name | Gene name |
|---|---|---|---|---|
| *Atp1b2* | *Sash3* | *Dnase1L1* | *Atrx* | *Tenm1* |
| *Itgb8* | *Utp14a* | *Slc10a3* | *Magt1* | *Ocrl* |
| *Prkcd* | *Elf4* | *Ubl4a* | *Atp7a* | *Phf8* |
| *Gjb6* | *Aifm1* | *Ikbkg* | *Taf9b* | *Huwe1* |
| *Pou6f1* | *Zfp280c* | *Tbl1x* | *Brwd3* | *Smc1a* |
| *Cdc25c* | *Enox2* | *Prkx* | *Chm* | *Klf8* |
| *Adnp2* | *Firre* | *Prrg1* | *Diaph2* | *Acot9* |
| *Uty* | *Mbnl3* | *Dmd* | *Pcdh19* | *Sms* |
| *Kcnt1* | *Hs6st2* | *Il1rapl1* | *Cstf2* | *Cnksr2* |
| *Kcnd3* | *Usp26* | *Pola1* | *Arl13a* | *Bclaf3* |
| *Prkag2* | *Gpc4* | *AL646049.1* | *Nox1* | *Sh3kbp1* |
| *Asb4* | *Plac1* | *Heph* | *Cenpi* | *Map3k15* |
| *Cav2* | *Fam122b* | *Ophn1* | *Drp2* | *Adgrg2* |
| *Fbln2* | *Fam122c* | *Stard8* | *Armcx4* | *Phka2* |
| *Rgma* | *Zfp36l3* | *Efnb1* | *Gprasp1* | *Cdkl5* |
| *Bckdk* | *Ints6l* | *Eda* | *Bhlhb9* | *Gja6* |
| *Ddx3y* | *Mmgt1* | *Awat2* | *Tbc1d8b* | *Nhs* |
| *Usp9y* | *Slc9a6* | *Otud6a* | *Rbm41* | *Piga* |
| *Shroom4* | *Fhl1* | *Igbp1* | *Tex13b* | *Gpm6b* |
| *Clcn5* | *Arhgef6* | *Kif4* | *Nxt2* | *Ofd1* |
| *Usp27x* | *Fgf13* | *Dlg3* | *Acsl4* | *Mid1* |
| *Foxp3* | *Tmem185a* | *Snx12* | *Tmem164* | |
| *Gpkow* | *Mamld1* | *Foxo4* | *Ammecr1* | |
| *Ccdc120* | *Gabre* | *Il2rg* | *Pak3* | |
| *Otud5* | *Zfp275* | *Zmym3* | *Lhfpl1* | |
| *Pcsk1n* | *Atp2b3* | *Ogt* | *Lrch2* | |
| *Gata1* | *Dusp9* | *Nhsl2* | *Alas2* | |
| *Fthl17f* | *Pnck* | *Rps4x* | *Apex2* | |
| *Rpgr* | *Slc6a8* | *Cited1* | *Tro* | |
| *Tspan7* | *Renbp* | *Dmrtc1b* | *Gnl3l* | |
| *Bcor* | *Mecp2* | *Zdhhc15* | *Fam120c* | |
| *Med14* | *Rbm10* | *Klhl13* | *Upf3b* | |
| *Nyx* | *Cdk16* | *Dock11* | *Tmem255a* | |
| *Cask* | *Usp11* | *Il13ra1* | *Lamp2* | |
| *Fundc1* | *Araf* | *Lonrf3* | *Cul4b* | |
| *Slc9a7* | *Elk1* | *Septin6* | *Gria3* | |

**Supplementary Table S2:** Specific force of EDL and soleus muscle from WT and dHT mice treated with vehicle or TMP269+5-Aza for 15 weeks. Muscles were stimulated with a single twitch or tetanic stimulation (EDL: 150 Hz, 400 ms duration; soleus 120 Hz, 400 ms duration). Values are expressed as specific force ($mN/mm^2$) *p <0.05 **p<0.01 dHT vs WT; ¶ p<0.05 dHT vehicle vs dHT TMP269+5-Aza (ANOVA followed by the Bonferroni post hoc test). (ANOVA followed by the Bonferroni post hoc test).

| | | EDL | | soleus | |
|---|---|---|---|---|---|
| **Genotype** | **Treatment** | **Twitch** (mean±SD) | **Tetanus 150 Hz** (mean±SD) | **Twitch** (mean±SD) | **Tetanus 120 Hz** (mean±SD) |
| WT | Vehicle (n=8) | 171.24±29.32 | 452.96±89.58 | 96.58±25.78 | 315.86±56.96 |
| dHT | Vehicle (n=10) (P value) | **64.92±13.93 (P=0.0015) | *373.76±73.16 (P=0.036) | *67.55±11.26 (P=0.040) | *276.29±40.04 (P=0.043) |
| | TMP269 + 5Aza (n=13) (P value) | *74.37±41.30 (P=0.012) | *379.96±87.87 (P=0.044) | *¶84.61±14.06 (*P=0.048) (¶P=0.047) | *¶334.78±65.74 (*P=0.047) (¶P=0.048) |

**Supplementary Table S3.** Fiber type composition of soleus muscles from mice treated for 15 weeks with vehicle or TMP269+5-Aza.

| | **MyHC I/total** (Ratio ±SEM) | **MyHC IIa/total** (Ratio ±SEM) | **MyHC IIx/total** (Ratio ±SEM) | **Total N° fibers counted** |
|---|---|---|---|---|
| **WT vehicle** | 0.31± 0.005 | 0.50±0.019 | 0.16±0.008 | 2881 |
| **dHT vehicle** | 0.33±0.02 | 0.53±0.02 | 0.13±0.008 | 2642 |
| **dHT treated** | 0.27±0.0059 | 0.56±0.015 | 0.15±0.01 | 2983 |

**Supplementary Table S4:** Analysis of electrically evoked calcium transients in single FDB muscle fibers isolated from WT and dHT littermates, treated with vehicle or TMP269+5-Aza (25 mg/Kg) for 15 weeks. *p<0.05 dHT vs WT;¶ p<0.05 dHT vehicle vs dHT TMP269+5-Aza (ANOVA followed by the Bonferroni post hoc test).

| | | **Treatment** | **Number of mice/N° of fibers analyzed** | **ΔF/F (mean±S.D.)** | **TTP msec (mean±S.D.)** | **HTTP ms (mean±S.D.)** | **HRT ms (mean±S.D.)** |
|---|---|---|---|---|---|---|---|
| **Twitch** | WT | Vehicle (NMP/PEG) | 4 (n=91) | 1.38±0.30 | 1.29±0.90 | 0.80±0.27 | 2.06±1.75 |
| | dHT | Vehicle (NMP/PEG) (P value) | 6 (n=110) | *0.98±0.22 (P=0.038) | 1.29±0.65 | 0.78±0.27 | 2.07±1.20 |
| | dHT | TMP269 +5Aza (P value) | 5 (n=155) | ¶1.21±0.28 (P=0.043) | 1.29±0.30 | 0.81±0.22 | 1.94±1.67 |
| **Tetanic** | WT | Vehicle (NMP/PEG) | 4 (n=63) | 1. 62±0.21 | | | |
| | dHT | Vehicle (NMP/PEG) (P value) | 6 (n=92) | *1.22±0.22 (P=0.039) | | | |
| | dHT | TMP269 +5Aza (P value) | 5 (n=78) | ¶1.42±0.23 (P=0.042) | | | |

**Supplementary Table S5:** Sequence of primers used and gene targets

| Primer details | | Primer sequence | |
|---|---|---|---|
| | | **Forward** | **Reverse** |
| Ex36 Ryr1 | Genotyping | TGCTGGCTTCAGAGTGATCG | CGAGGGAAGTTGAGGTTGGG |
| Ex91 Ryr1 | Genotyping | GAGATGTTCGTGAGTTTCTGCGAGG | TGAGGGTTGTTCTTGGTGTATTTGG |
| Ryr1 | qPCR | CGCCAAAACGGAGAGAAAGTC | TTGATGGTGGTCGTGTTCCC |
| Cacna1s | qPCR | TCAGCATCGTGGAATGGAAAC | GTTCAGAGTGTTGTTGTC |
| Hdac4 | qPCR | CACTGCATTTCCAGCGATCC | AAGACGGGGTGGTTGTAGGA |
| Gapdh | qPCR | CTGCACCACCAACTGCTTAGC | GGCATGGACTGTGGTCATGAG |

**<u>Supplementary Table S6</u>:** List of antibodies and suppliers

| **Target** | **Supplier** | **Catalog number** |
|---|---|---|
| Acetyl Lysine | Abcam | Ab21623 |
| Alexa Fluor 568 conjugated. Mouse IgG1 | ThermoFisher Scientific | A21124 |
| Alexa Fluor 488 conjugated. Mouse IgM | ThermoFisher Scientific | 21042 |
| Alexa Fluor 647 conjugated. Rabbit IgG | Jackson ImmunoResearch | 711-605-152 |
| Calsequestrin 1 | Sigma | C0742 |
| Cav1.1 | DSHB | IIC12D4 |
| DyLight 405 Alexa Fluor 488 conjugated. Mouse IgG Fcγ2b | Jackson ImmunoResearch | 115-475-207 |
| Histone H3 | Abcam | Ab1791 |
| Histone H3 (acetyl K9) | Abcam | Ab10812 |
| JP-45 | Made in-house | (Zorzato et al. 2000)* |
| Laminin | Sigma | L9393 |
| MyHC | Santa Cruz | Sc-376157 |
| MyHC I | DSHB | BA-D5 |
| MyHC-IIa | DSHB | SC-71 |
| MyHC IIb | DSHB | BF-F3 |
| RyR1 | Cell Signaling | D4E1 |
| SERCA1 | Santa Cruz | sc-8093 |
| SERCA2 | Santa Cruz | sc-8095 |

* F Zorzato, A Anderson, K Ohlendieck, G Froemming, R Guerrini, S Treves. Identification of a novel 45 kDa protein (JP-45) from rabbit sarcoplasmic reticulum junctional face membrane. *Biochem. J.* **351,** 537-543 (2000).
